# Global Insights into Evolution and Adaptation in KPC and NDM Coproducing Carbapenem-Resistant *Klebsiella pneumoniae*

**DOI:** 10.1101/2025.10.16.682762

**Authors:** Mingxiao Chen, Xiwei Zhang, Jingyi Zhang, Yu Gan, Zhuoyan Zhong, Tingting Deng, Yitong Han, Jingjie Li, Xiaobing Duan, Jingjie Song, Qiang Zhou

**Author notes:** Correspondence: Qiang Zhou; Jingjie Song; Xiaobing Duan. These authors contributed equally to this work (Mingxiao Chen, Xiwei Zhang, Jingyi Zhang).

## Abstract

**Objectives:** The worldwide dissemination of carbapenem-resistant *Klebsiella pneumoniae* co-producing KPC and NDM (KPC-NDM-CRKP) poses a substantial clinical challenge.

**Methods:** This study characterized 5 KPC-NDM-CRKP isolates from a tertiary hospital in southern China and integrated 460 global isolates genomes to demonstrate the population framework of this epidemic pathogen.

**Results:** 5 clinical isolates were multidrug resistance, exhibiting high level resistance to first-line drugs including carbapenems and ceftazidime/avibactam. Globally, KPC-2-NDM-1 (50.3%) were the most common and an increasing trend of ST11-KL64 (20%) isolates carrying *iucA* and *rmpA*/*A2*. Resistance and virulence genes were positively associated with specific mobile elements including *bla*_KPC-3_ with Tn*4401, iucABC* with IncHI1B, *iroBCDN* with IncQ1. The phylogenetic analysis revealed distinct lineage-specific mutations and pan-genome enrichments synergistically promoted adaptive evolution: cluster 1 had membrane transport and secretion but enhanced immune evasion and horizontal gene transfer, cluster 3 adapted metabolically for nutrient-limited niches; cluster 4 reprogrammed metabolism for host adaptation, cluster 6 displayed membrane repair and reduced virulence (e.g., KC187). High-risk *iucA/rmpA*-positive isolates in cluster 2 acquired enhanced virulence and metabolic adaptive capacity through recombination hotspots (ABC transporters DdpABCD, efflux systems MdtABCD, metabolic enzymes XylB and MtlD, regulators BaeR and OmpR) and pan-genome enrichment (siderophore biosynthesis, toxin–antitoxin VapC and restriction-modification systems). Functionally, these traits correlated with high mucoviscosity, serum resistance, siderophore production and lower survival rates in *G. mellonella* that contribute to epidemiological success.

**Conclusions:** These findings demonstrated the discrete mutation, horizontal transfer, and accessory genome plasticity collectively drove the epidemiological success of KPC-NDM-CRKP.

## Introduction

Carbapenem-resistant *Klebsiella pneumoniae* (CRKP) has emerged as a critical global public health threat, primarily due to its ability to resist multiple classes of antibiotics, severely limiting therapeutic options and contributing to high mortality rates among immunocompromised patients(1, 2). The primary mechanism driving CRKP resistance is the production of carbapenemases, a highly diverse class of β-lactamases. *Klebsiella pneumoniae* carbapenemase (KPC) encoded by *bla*_KPC_ gene and New Delhi metallo-β-lactamase (NDM) encoded by *bla*_NDM_ gene are the most clinically relevant carbapenemases in CRKP(3, 4). The emergence of CRKP strains co-producing KPC and NDM (KPC-NDM-CRKP) represents an exceptionally grave threat (5).

The epidemiology of KPC-NDM-CRKP is predominantly facilitated by horizontal gene transfer, including *bla*_KPC_-positive strains acquiring *bla*_NDM_-carrying plasmids(5). The acquisition of resistance determinants underscores the remarkable adaptability of *K. pneumoniae*, facilitating the emergence of highly resistant strains under selective antibiotic pressure(6). The emergence of hypervirulent carbapenem-resistant *Klebsiella pneumoniae* (hv-CRKP) poses a critical public health threat due to its dual capacity for high pathogenicity and resistance to first-line antimicrobial agents(7, 8). At present, the report of hv-CRKP co-producing NDM and KPC is rare and there remain unclear on the global spread and evolution patterns of KPC-NDM-CRKP. In this study, we described 5 KPC-NDM-CRKP isolates identified from senior patients in a tertiary hospital in south China, which exhibited different sequence types and capsules (ST11-KL64, ST15-KL19 and ST1869-KL2). We analyzed the characterization and molecular mechanisms of multidrug resistance and hypervirulence among KPC-NDM-CRKP using whole-genome sequencing. Combining the available public database, a genomic epidemiological investigation was conducted to elucidate the population structure and genomic framework of this significant bacterial pathogen.

## Method

### 2.1 Bacterial collection

KPC-NDM-CRKP isolates KC141, KC187, KC258, KC263 and KC433 were collected from patients in the Intensive Care Unit of the Second Affiliated Hospital of Guangzhou Medical University from March to October 2024. Both the information of patients and 5 isolates had obtained ethical permission. A total of 458 KPC-NDM-CPKP isolates were retrieved from the NCBI GenBank (https://www.ncbi.nlm.nih.gov/pathogens/) as of August 12, 2024, and the geographical location, year and accession number were listed in Table S1.

### 2.2 Antimicrobial susceptibility testing

The MIC values of 5 KPC-NDM-CRKP isolates for various antimicrobial agents were performed using VITEK-2 Compact equipment (Table S2). The carbapenemase production was examined using both the modified carbapenem inactivation method (mCIM)/EDTA-modified CIM (eCIM) and APB/EDTA methods. The interpretation of the results was conducted by the Clinical and Laboratory Standards Institute (CLSI M100) guidelines(9).

### 2.3 String test

The strains were inoculated onto agar media and aerobically incubated at 35 °C for 20– 24 h. The colonies on the agar media were pulled up using a 10µL plastic loop, and the viscous string formation was observed. A positive string test result was defined as a viscous string formation of >5 mm at one or more spots.

### 2.4 Quantitative siderophore production assay

To access the production of bacterial siderophore, Kings B agar plates containing chrome azurol S dye (CAS) were prepared as described by Tian et al (10). Specifically, 10 mL of stationary-phase, iron-chelated cultures were deposited on CAS plates at 37°C. After 48 hours of growth, the formation of an opaque golden-yellow zone around the colony was used to identify high-level siderophore production.

### 2.5 Whole-genome sequencing and bioinformatics analysis

KC258 and KC433 were sequenced with Illumina Novaseq PE150 platform. KC141, KC187 and KC263 were long-read sequenced with PacBio Sequel platform. genomes were assembled by Unicycler v0.5.0 and annotated by Prokka v1.14.5 (11, 12). All genomes were then used to construct the pangenome with Panaroo v1.2.9(13). Kleborate v2.2.0 was used to determine sequence types, capsule types, virulence genes and antimicrobial resistance genes (ARGs) and VGs (virulence genes)(14). Mobile genetic elements (MGEs) were detected by bacant(15). The full of 465 KPC-NDM-CRKP genomes was used to generate a core-genome single nucleotide polymorphism (SNP) using Snippy v.4.1.0 and to recognize recombination-free sequence using Gubbins v.2.2.0(16). A maximum likelihood (ML) phylogenetic tree was constructed by IQ-Tree2 v.1.6.7.2(17). All genomes were annotated and then used to construct the pangenome with Roary and Panaroo. RhierBAPS was used to detect Bayesian structures (18). We aligned genomes in the same cluster using EToKi align and identified the recombination using RecHMM. High-recombination region (HRR) was defined as continuous regions with >100 recombination events in a sliding window of 100Kb.

### 2.6 Conjugation assay

5 KPC-NDM-CRKP iaolates served as donor strain and rifampicin-resistant *E. coli* C600 was used as the recipient strain. Transconjugants were selected on Luria-Bertani plates containing 100 µg/mL rifampicin and 2 µg/mL imipenem, and the resistance phenotypes were determined using the VITEK-2 Compact equipment.

### 2.7 G. mellonella in vivo infection model

We also tested the virulence of 5 KPC-NDM-CRKP isolates by *G. mellonella* infection assays. Overnight cultures of *K. pneumoniae* strains were resuspended in PBS to achieve final concentrations of 1×10^6^ CFU/mL. These *G. mellonella* were injected 10µL of the bacterial suspension and maintained at 37°C. The survival rate was calculated after 36h of incubation.

### 2.8 Statistics

Statistical significance was assessed using a two-tailed Student’s t-test and log-rank (Mantel-Cox) test of the GraphPad Prism 9 software. P<0.05 was regarded as statistically significant. A log-rank (Mantel–Cox) test was performed for the survival curves. ****P < 0.0001.

## Results

### 3.1 The antimicrobial resistance and virulence-related features of 5 clinical KPC-NDM-CRKP isolates

Patients with KPC-NDM-CR-KP infections were admitted to a tertiary hospital’s ICU in Guangzhou from March to October 2024. Infection sites comprised pneumonia (2/5), urinary tract (1/5), ear canal (1/5) and biliary tract infection (1/5) (Table S1). Most patients had favorable clinical outcomes, whereas patients with KC187 and KC263 had poorer prognoses due to severe pneumonia and ear canal infection respectively. Among 5 KPC-NDM-CRKP isolates, KC187 was ST15-KL19, KC263 belonged to ST1869-KL2 and remaining isolates were ST11-KL64 (Table S2). These carbapenem-resistant isolates were multidrug-resistant phenotypes, which also showed resistance to β-lactam, β-lactamase inhibitors, quinolone and sulfonamides but susceptible to colistin. The mCIM results were interpreted as carbapenemase positive. For the APB/EDTA results, it showed the co-production of class A carbapenemase and MBL.

The representative ST-KL types of isolates KC141, KC258 and KC263 were selected to demonstrate the genetic environment of *bla*_KPC-2_ and *bla*_NDM-5_. *bla*_KPC-2_ gene located on non-mobilizable IncR plasmids (KC141 and KC263) and chromosome of KC187, with surroundings of IS*26*-*Tn3*-IS*Kpn27*-*bla*_KPC-2_-IS*Kpn6* that was different from classical Tn*1721* transposon (Fig. 1A and B, Table S3). *bla*_NDM-5_ was carried by IS5D-*bla*_NDM-5_-*ble*_MBL_-IS*26* and located on conjugative IncX3 plasmids harboring type IV secretion system (T4SS) for horizontal transfer (Fig. 1C). Conjugation experiments revealed that *bla*_NDM_-positve plasmids could be transferred to the recipient *E. coli* C600 (Table S4).

**Figure 1.**
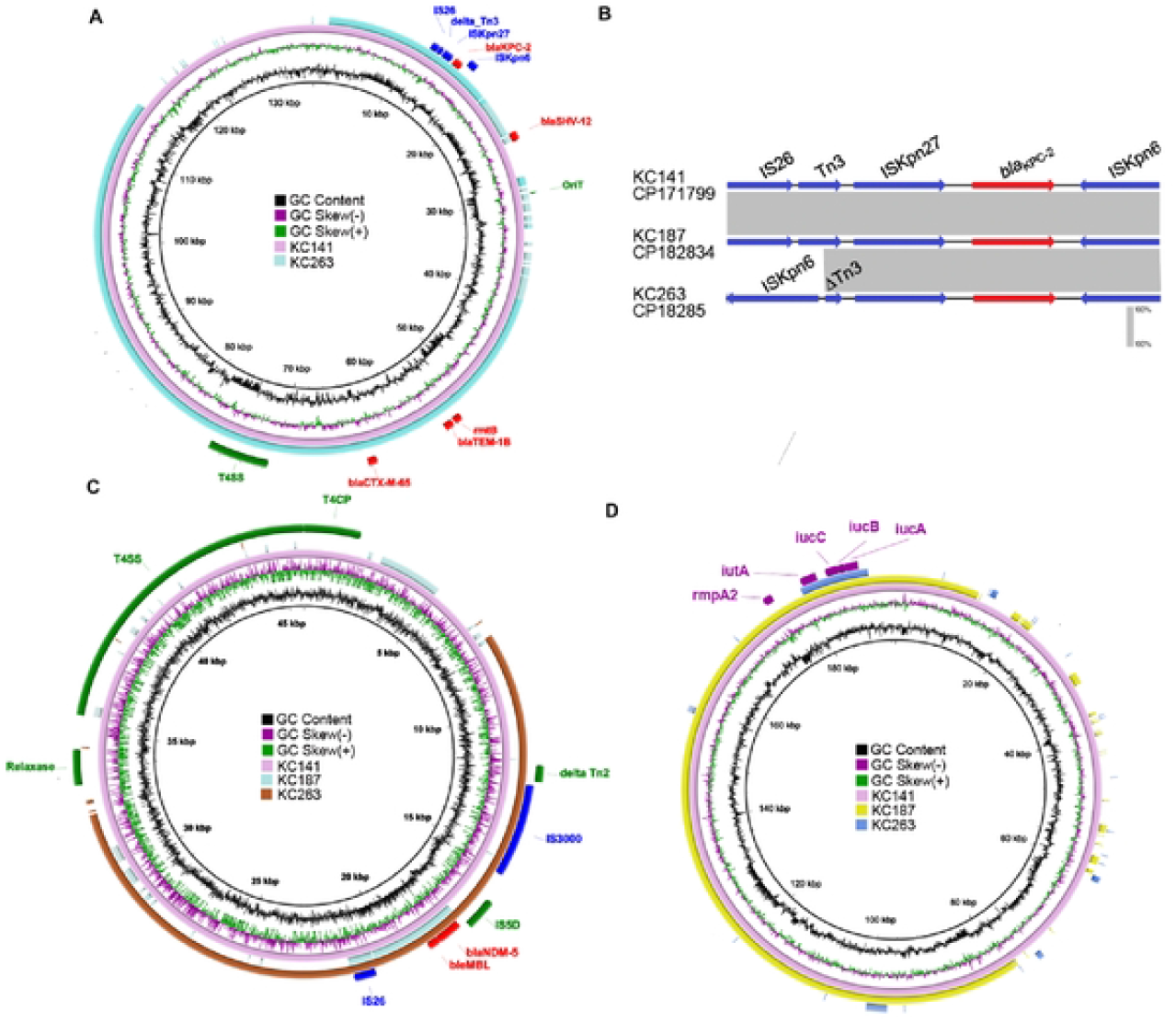
Comparative analysis of *bla*_KPC_-positivc, *bla*_NDM_-positive and virulence plasmids in KPC-NDM-CRKPs. (A) Genome alignment was performed,with *bla*_KPC-2_-positive plasmids. (B) Comparison of the genetic environment of *bla*_*KPC-2*_gene harboring in plasmids or chromosome. (C) Genome alignment was performed with *bla*_NDM-5_-positive plasmids. (D) Genome alignment was performed with virulence plasmids.

### 3.2 Overview of global 465 KPC-NDM-CRKP isolates

To obtain a comprehensive overview of NDM-KPC-CRKP, 5 isolates in this study were integrated with additional 460 public NDM and KPC co-producing genomes to form a global collection of 465 isolates. These isolates were frequently identified in Asia (47.1%), followed by North America (29.7%) and Europe (12.3%) (Fig. 2A and B). For carbapenemase gene phenotype, the most prevalent was KPC-2-NDM-1 accounting for 50.3%, followed by KPC-2-NDM-5 (18.3%) and KPC-3-NDM-1 (13.3%) (Fig. 2C). Several KPC-NDM types are frequently associated with geographic locations, comprising KPC-2-NDM-5 in China (47.9%), KPC-3-NDM-5 (81.2%) and KPC-3-NDM-7 (100%) in the United States (Table S5).

**Figure 2.**
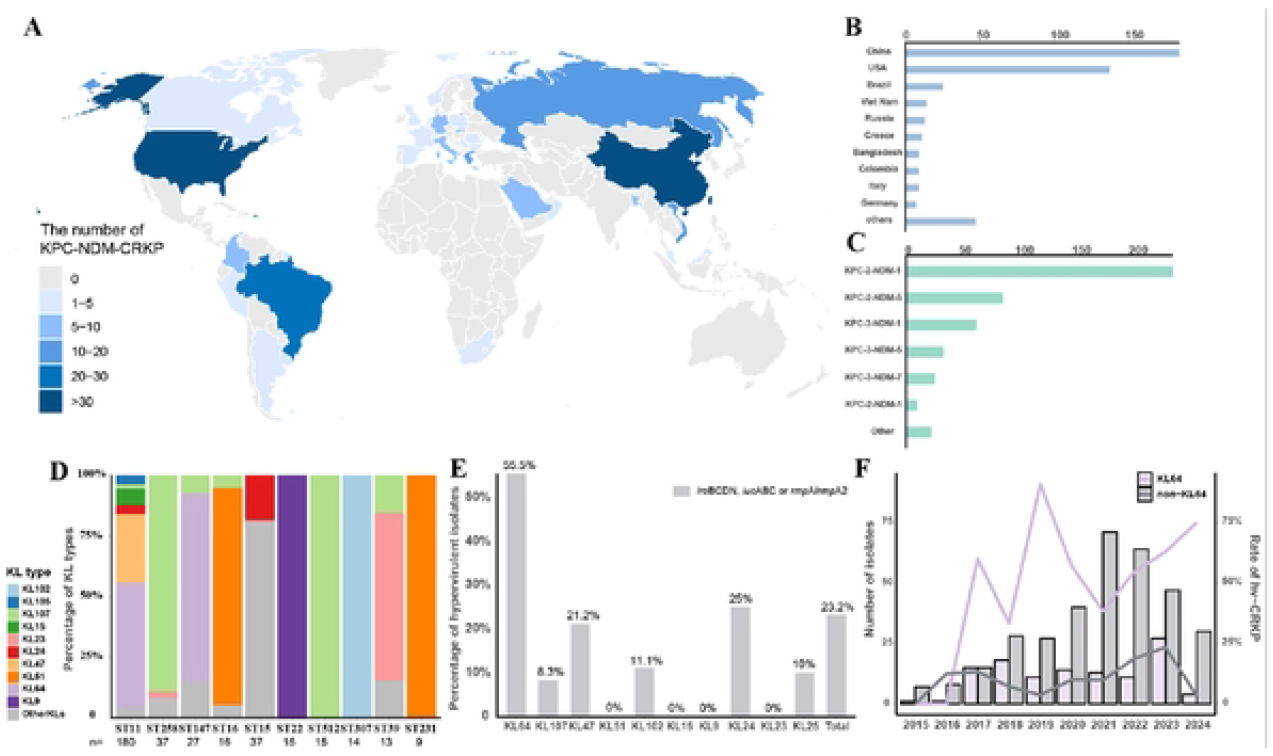
Geographic distribution and genetic framework of 465 KPC-NDM-CPKP isolates. (A) The map shows the distribution of all isolates across continents. (B) Bar graph of the number of isolates by different countries. (C) Bar graph of the number of all isolates by different carbapcncmasc genes subtypes. (D) Percentage of K types between different STs. (E) The proportion of hv-CRKP in different K-types CRKP isolates. The grey bars indicate strains,with hypervirulent genes (*iroBCDN, iucABC* or *rmpA*/*rmpA2*). (F) Trends in the proportion of CRKPs from KL64 and non-KL64. The bar represents the number of CRKP in different K types. The line represents the percentage of CRKP carrying virulence genes (*iroBCDN, iucABC* or *rmpA*/*rmpA2*) in different K types.

MLST assigned the 471 isolates to 64 distinct STs, with the majority being ST11 (180/465, 38.7%). KL64 was the predominant KL type in ST11 and ST147, accounting for 52.0% (93/179) and 77.8% (21/27) respectively. Notably, KL64 (55.5%) had significantly higher number of hypervirulent genes (*iucA* and *rmpA*/*rmpA2*) than the non-KL64 population, including KL24 (25%) and KL47 (21.2%) (Figure 3E). The population of non-KL64 hv-CRKP showed a fluctuating expansion trend between 2015 and 2023, followed by a sharp decline (Fig. 2D).

**Figure 3.**
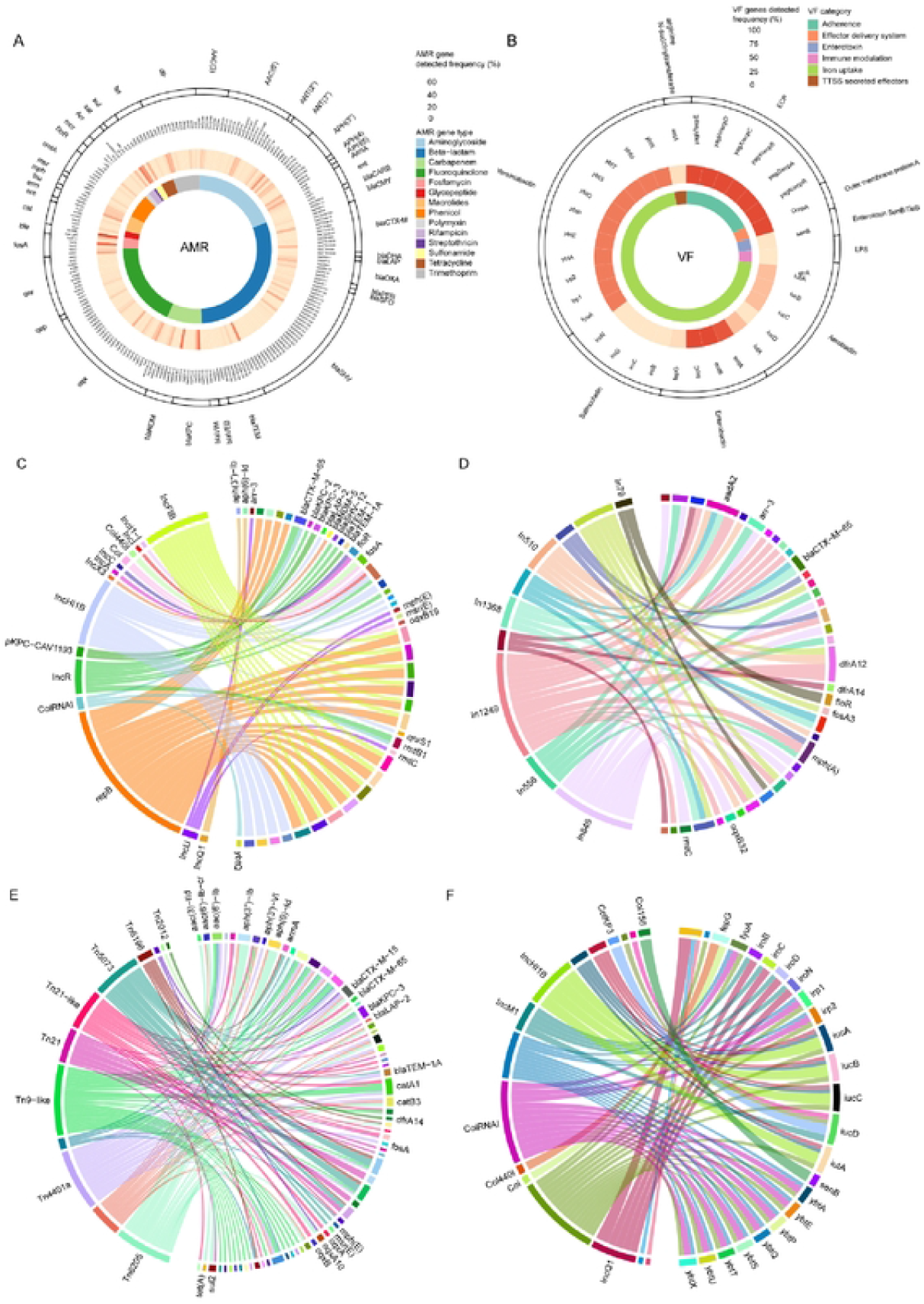
AMR and VF potential and their associated mobile elements among the KPC-NDM CRKP. (A) The frequencies of AMR genes identified in KPC-NDM-KP. (B) The frequencies of VF genes identified in KPC-NDM-KP. (C) Relationship between the major ARGs and plasmid Inc types. (D) The relationship between the major ARGs and integron vehicles. (E) The relationship between the major ARGs and transposon vehicles. (F) Relationship between the major VGs genes and their plasmid Inc types.

### 3.5 Genome relationship between major ARGs, VGs and MGEs

465 NDM-KPC-CRKP isolates harbored a wide variety of ARGs, VGs and MGEs. A total of 248 different ARGs were identified, with the majority being carbapenemase genes *ble*_MBL_ (n=435), *bla*_KPC-2_ (338) and *bla*_NDM-1_ (298) (Fig. 3 A and B, Table S5). Of note, most isolates carrying carbapenemase genes were also positive for *fosA6* (333), *bla*_TEM-1_ (275) and *bla*_SHV-158_ (164). An association study between ARGs and genetic mobile elements was as follows: (I) plasmids: IncR with *bla*_CTX-M-65_ and rmtB1; pKPC-CAV1193 with *bla*_KPC-3_. (II) Ins: In*1249* with *aadA2* and *dfrA12*; In*556* with *arr-3*; In*1368* with *bla*_CTX-M-65_. (III) Tns: Tn*6205* with *sul2, bla*_OXA-1_ and *tet(A)*; Tn*4401* with *bla*_KPC-3_ and *bla*_TEM-1A_; Tn*21*-like with *oqxAB* and *catA1* (R>0.3, p<0.01) (Fig. 3 C, D and E, Table S6).

All isolates harbored a diverse set of virulent genes (VGs), *entA, entB, fepC* and *ecpABCDER* that enhanced the pathogenicity were detected in over 90% of isolates. Hypervirulent genes *ybt* (335/465, 72.0%), *iutA-iucABC* (131/465, 28.2%), iro*BCDN* and *rmpA*/*rmpA2* (108/465, 23.2%) genes were also found in many isolates. Moreover, VFs show a coexistence relationship with specific plasmids: IncQ1 with *iroBCDN*; IncHI1B with *iutA-iucABC*; ColKP3 and IncM1with *iucD*; ColRNAI with *ybtAEPQSTUX* (R>0.3, p<0.01) (Fig. 3F).

### 3.6 phylogenetic tree and pan-genome analysis of global KPC-NDM-CRKP

The phylogenetic tree ranging from 0 to 131354 SNPs distances can be divided into seven clusters and was mainly composed of cluster 2 (189/465, 40.6%) and cluster 5 (98/465, 21.1%) (Fig. 4A). Four isolates (KC141, 258, 263 and 433) recognized from this study gathered in cluster 2, while KC187 was classified into cluster 6. Cluster 2 predominantly comprised KPC-2-NDM-1 (113/189, 59.8%) and KPC-2-NDM-5(65/189, 34.4%). Cluster 5 was dominated by KPC-2-NDM-1 (41/98, 41.8%) and KPC-3-NDM-7 (20/98, 20.4%). Notably, isolates in cluster 1 showed missense mutations in the *mdtK* (g.207C>G), *mdtH* (g.388A>G and 374T>A) and *mdtM* (g.226A>G) genes, whereas those isolates in cluster 2 had missense mutations in *ompA* gene (g.215G>A, 205A>C and 180A>G).

**Figure 4.**
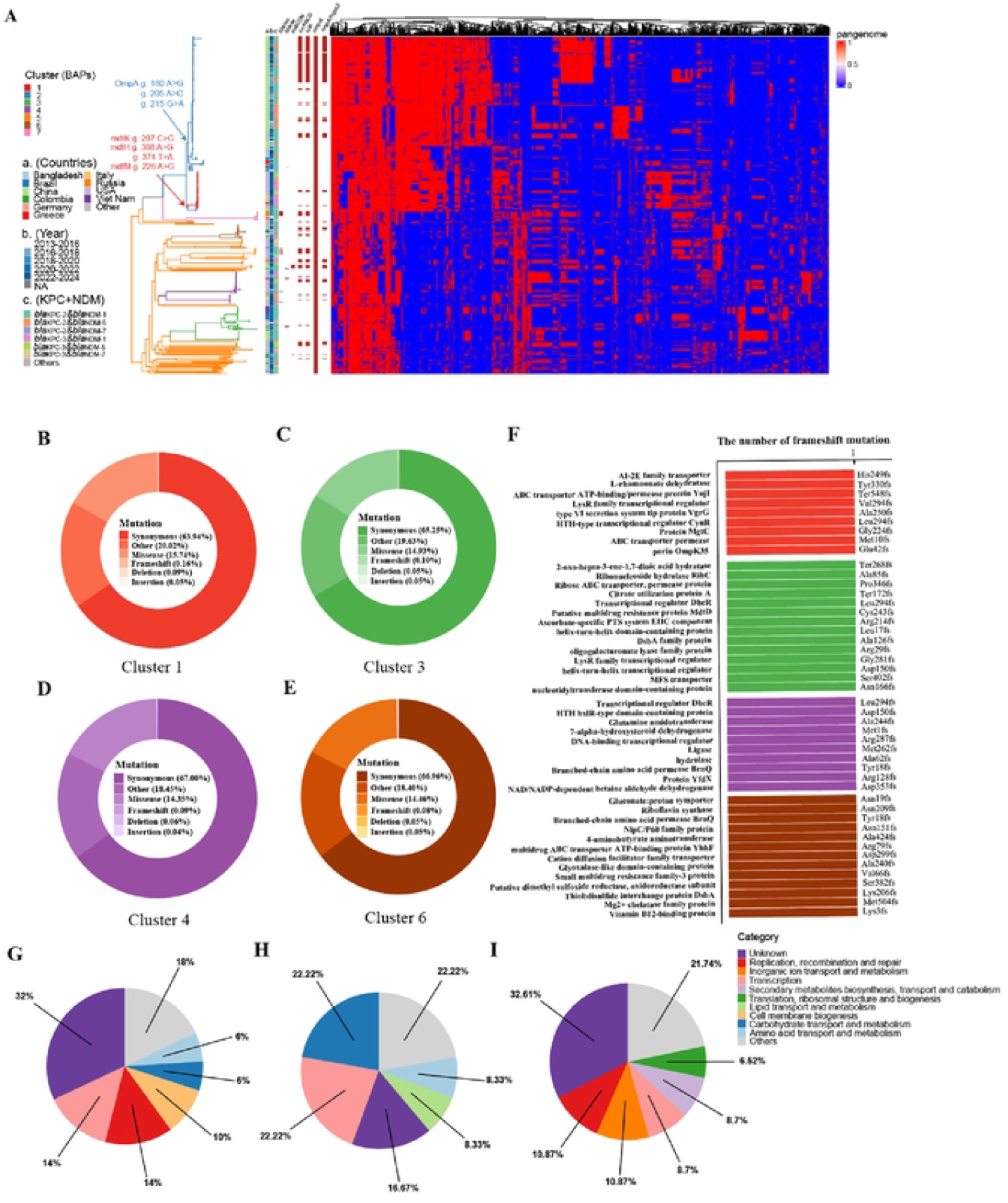
Phylogenetic analysis and pan genome contents of KPC-NDM-CRKP. (A)The branch colors are denoted by BAPs of population. Annotations from the inner to the outer rows include countries, year and KPC-NDM subtypes. Additional details regarding SNPs of all isolates can be found in Table S4. (B-E) The type and proportion of frameshift mutations in different clusters. The depth color of the circle graph represents different types of mutation. (F) The bar chart showed the distinct frameshift mutations in gene products among each cluster. (G-H) Functional categories of unique pangenome in cluster I compared,with cluster 2 (G), in cluster 2 compared with cluster I (H) and in *iucA*-positive isolates of cluster 2 compared with iucA-negative isolates in cluster 2 (I).

We further investigated how functional genes with frameshift mutations shaped the evolution of different clusters. Cluster 1 exhibited in membrane transport (OmpK35) and secretion systems, (VgrG, MgtC) and regulators (LysR and CynR families). Cluster 3 displayed in carbohydrate and nucleotide metabolism (RihC, citrate utilization protein A, oligogalacturonate lyase) and transport (MFS, MdtD) (Fig. 4B). Cluster 4 related to amino acid and redox metabolism genes (glutamine amidotransferase, 7α-hydroxysteroid dehydrogenase, betaine aldehyde dehydrogenase), together with transcriptional regulators (DhcR, HTH hxlR-type) and stress-response factors (YfdX) (Fig. 4C). Cluster 6 affected transport (gluconate/proton symporter, BrnQ, cation diffusion facilitator, multidrug transporters), metabolism and cofactors (riboflavin synthase, 4-aminobutyrate aminotransferase, Mg^2+^chelatase, vitamin B12-binding protein) and stress response (DsbA, NlpC/P60 family protein) (Fig. 4D).

The results of pangenome analysis in ML tree identified a total of 19,509 genes among 465 genomes and 16,604 accessory genes. We further focus on the accessory genome of major clusters (I and II) in different regions (Table S7). In a comparative analysis between cluster 1 and cluster 2 (genes present in >90% of subject cluster and in <10% of control cluster). Cluster 1 characteristically harbored gene sets correlating to capsule biosynthesis (*wzi*), cell membrane synthesis (*yibD, rfbN*), prophage integrases (*alpA*) and plasmid conjugal transfer (*traD, MobA*/*MobL* family) (Fig. 4G). Cluster 2 characteristically carried gene involved in carbohydrate metabolism (*puuE, ggt1, gatA, guaA, fabF, pgm, celB, cbhA*) and transcriptional regulators (*mobC*, AraC-like domain, helix-turn-helix proteins) (Fig. 4H).

### 3.7 The recombination and hypervirulence features of *iucA*-positive isolates in cluster 2

The recombined region in *iucA*-positive population in cluster 2 contained genes related to ABC-type systems (*ddpABCD, gsiA*), multidrug efflux pumps (*mdtABCD*), metabolic enzymes (*xylB, mtlD, dcd, udk*), regulators (*baeR, baeS, ompR, kdpD, csgD*) and membrane-associated proteins (*asmA, ccdA, yegD, yegS, yegQ*) (Fig. 5A and B). In addition, these isolates acquired the stop-lost mutation (cytoplasmic alpha-amylase) and missense mutations (2-alkyl-3-oxoalkanoate reductase, bifunctional aspartokinase/homoserine dehydrogenase, the replication initiation protein) that involve carbohydrate metabolism, amino acid synthesis and DNA replication (Fig. 5C). Notably, accessory genomes of *iucA*-positive isolates were specially enriched in metabolic adaptation genes (*thrS, hemB, pyrC, def*, cobalamin biosynthesis) toxin– antitoxin components (*vapC*), restriction endonuclease, resolvase (pin) and several transposases (Fig. 4I).

**Figure 5.**
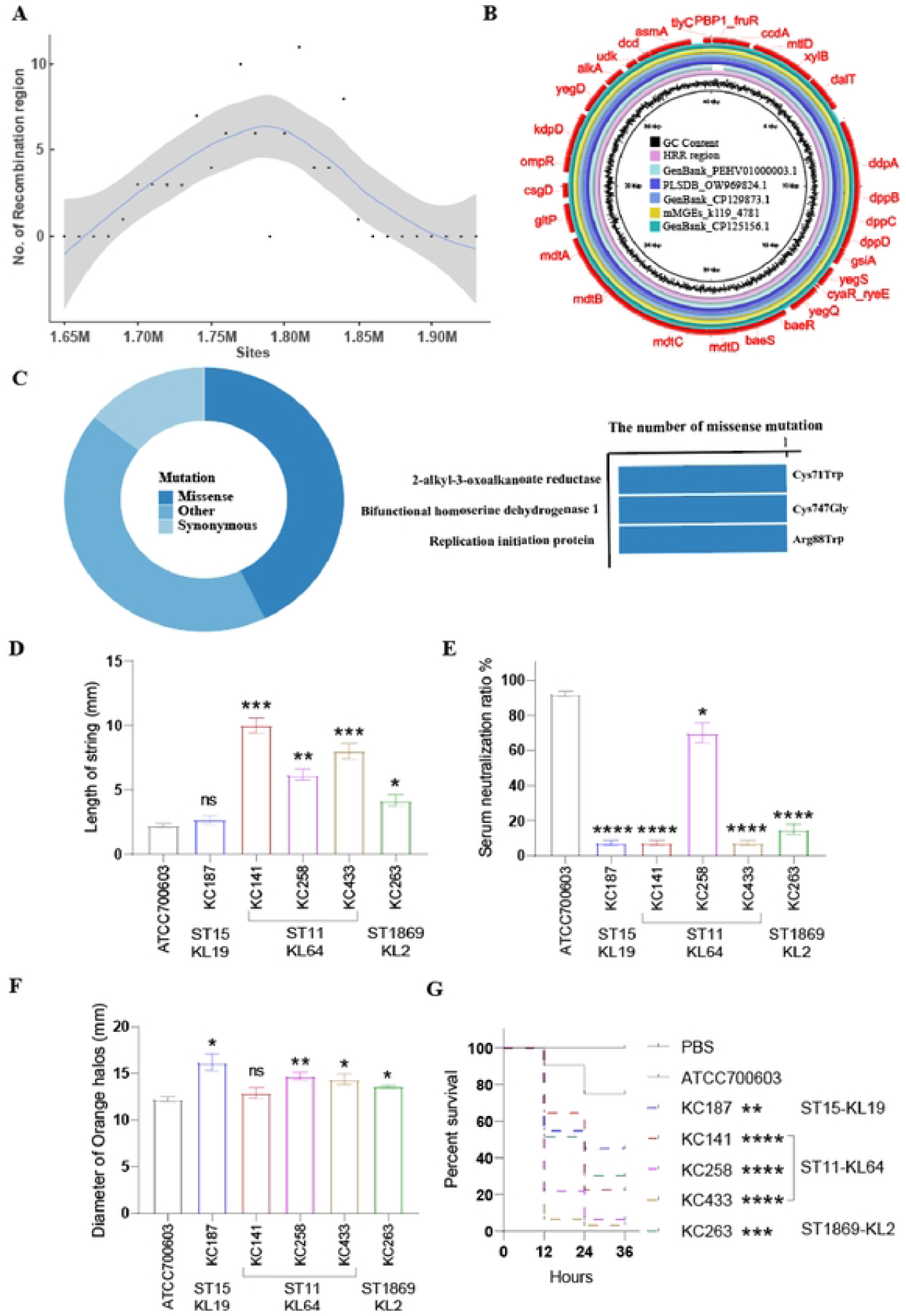
The recombination events and virulence phenotypes of *iucA*-positive population. (A) The number of recombination events of HRR in the chromosome. (B) Sequence comparison of HRR and plasmid vehicles,with high similarity. (C) The types and proportions of nucleotide mutations. (D) String test data. (E) Serum killing assay in vitro.(F) Siderophores production is determined by CAS agar plate. (G) The survival curves of *G. mellonella* infected by 5 clinical KPC-NDM-CRKP isolates and control strain (ATCC700603). ••••p <0.0001.

Four isolates (KC141, 258, 263 and 433) in *iucA*-positive population of cluster 2 were used for virulence evaluation. String test showed these isolates were high mucoviscosity except for KC187 in cluster 6 (Fig. 5D). All isolates had greater serum resistance than ATCC700603 (Fig. 5E). Quantitative siderophore assays indicated that most isolates produced more siderophores except for KC141 (Fig. 5F). 5 KPC-NDM-CRKP isolates (3.23∼45.16%) show greater pathogenicity than ATCC700603 (75.00%) in *G. mellonella* assay (Fig. 5G). Moreover, 4 *iucA*-positive isolates (3.23∼30.30%) displayed lower *G. mellonella* survival rate than KC187 (45.16%).

## Discussion

CRKP has been classified in the critical-priority tier by the WHO due to significant ability to acquire MGEs, which serve as vehicles for conferring both multidrug resistance and hypervirulence(19). In this study, 5 KPC-NDM-CRKP isolates were recovered from senior patients in the ICU of a tertiary hospital, which caused long-term colonization and high mortality rates (40%). These carbapenem-resistant isolates carried various ARGs and exhibited resistance to commonly used clinical antimicrobial agents, including β-lactams and β-lactams/β-lactamase inhibitor combinations that have been suggested as last-line drugs for treating CRKP infections(20). Genetic environmental analysis indicated *bla*_KPC-2_ was moved by Tn*1721* transposon and *bla*_NDM_ gene was moved by IS*26*, whose host plasmids display diverse sequences and their evolution contributed to the clonal spread of CRKP (21). The *bla*_NDM_-positive plasmids were conjugative that is consistent with previous research that the initial KPC-CRKP acquired a transferable *bla*_NDM_ plasmid (5, 22).

For 465 global isolates, ST11 is the dominant type of CRKP in China that evolved different clades with adaptive advantage (23). Notably, the proportion of ST11-KL64 carrying hypervirulent genes showed an increasing tendency, which represents a globally disseminated clade of CRKP that is known for hypervirulence and multidrug resistance(24). KPC-2-NDM-1 (50.3%) was the most common carbapenemase subtype in total 465 isolates, which could be due to steady stay in the environment and nosocomially transmission among patients(25). Many ARGs were positively associated with specific MGEs including *bla*_KPC-3_ with pKPC-CAV1193 (R>0.5); *bla*_CTX-M-65_ with IncR and In*1368*; *bla*_TEM-1A_ with Tn*4401*; *bla*_OXA-1_ with Tn*6205*, which were rarely observed in previous studies. For VFs profiles, *rmpA*/*rmpA2* and *iucABC*/*iutA*/*iroBCDN* that served as critical virulence determinants were present in more than 20% of all isolates, heightening the risk of transmission (26). These genes were strongly related to some plasmids (R>0.4) including *iroBCDN* with IncQ1; *iucD* with ColKP3 (R>0.6), which were rarely reported in previous publications. These findings underline that mobile elements play a crucial role in the capture and mobilization of ARGs and VGs.

The phylogenetic tree analysis revealed that isolates from cluster 1 showed multiple mutations in multidrug-resistant efflux pump genes *mdt*H, *mdt*K and *mdt*M that contributed to regulating antibiotic resistance including tetracycline, chloramphenicol(27, 28). These lineage-specific frameshift mutations underscore diverse adaptive strategies within CRKP evolution. Cluster 1 appears to prioritize envelope remodeling and attenuated secretion to promote antibiotic persistence and immune evasion(29-31). Cluster 3 reflects extensive disruptions in regulatory and metabolic genes that shaped genome plasticity in fluctuating environments(32, 33). Cluster 4 demonstrates loss of key biosynthetic and stress-response functions, indicating a transition toward host dependence and metabolic simplification(34, 35). Cluster 6 impairs nutrient uptake, multidrug efflux, cofactor biosynthesis and stress persistence that shaped adaptive fitness(36, 37). Pan-genome analysis revealed some clusters harbor distinct accessory genomes characterized by high plasticity. Cluster 1 appears to emphasize capsule variability, genome mobility, various conjugation elements and glycosyltransferases, which confer advantage on surviving in host immunity and consequent dissemination(38, 39). Clusters 2 carried *ompA* mutant and alternat metabolism of carbohydrate, amino acids and fatty acids that were associated with the bacterial adhesion and invasion(40, 41).

For *iucA*-positive population in cluster 2, the recombination region integrates Mdt transports, metabolic genes and regulators to facilitate membrane remodeling, energy flexibility, pathogenicity, drug and stress resistance(42-44). Especially OmpR enhances virulence and BaeSR two-component activates the expression of MdtABCD transporter to mediated multidrug resistance (45, 46). The observed SNP mutations involve pathways related to carbohydrate metabolism, amino acid synthesis and DNA replication. Additionally, the enriched pan genes of *iucA*-positive isolates corresponded to siderophore biosynthesis, toxins, antioxidation and ABC transporters that were vital for virulence and fitness(47, 48). ABC transporters participated in the biosynthesis of different classes of polysaccharides and antioxidation was linked to heightened virulence in ST11-KL64 hv-CPKP(49, 50). Together, these changes reflect adaptive metabolic capacity, genomic stability and ecological competitiveness of this high-risk population. The hypervirulent traits of high mucoviscosity, serum resistance, elevated siderophore production and higher *G. mellonella* mortality were presented in 4 representative *iucA*-positive isolates, highlighting the critical need for implementing routine surveillance(51, 52).

In conclusion, this study provides critical insights into the evolutionary dynamics and adaptive mechanisms driving the microevolution of KPC-NDM-CRKP. The distinct phylogenetic clusters employ varied adaptive strategies including membrane transport and repair, secretion systems and metabolic reprogramming, enabling niche specialization and host adaptation. We have traced the recombination hotspots, accessory genome plasticity, functional adaptations and enhanced pathogenicity of high-risk *iucA*-positive population to support the development of targeted interventions.

## Transparency declarations

The authors declare that they have no known competing financial interests

## Author contributions

Q.Z, J.S and X.D designed and supervised the study. M.C and J.Z performed the antimicrobial susceptibility tests and WGS. J.S and X.Z performed bioinformatics analysis. Y.G, Z.Z and T.D discussed the data and prepared the draft, figure and supplementary materials. Y.H and J.H collected the data and manuscript revision. X.D acquired the funding. All authors reviewed and approved the final manuscript.

## Data Availability

This project was funded by the National Natural Science Foundation of China (No.32470961), the Guangdong Basic and Applied Basic Research Foundation (No.2025A1515010102 and No.2024A1515012563).

## Ethical Declarations

The ethical approval number for this project is LYZX-2025-086-01.

